# *Ubc9* deletion in adipocytes causes lipoatrophy in mice

**DOI:** 10.1101/2020.09.12.294629

**Authors:** Aaron R. Cox, Natasha Chernis, Kang Ho Kim, Peter M. Masschelin, Pradip K. Saha, Shawn M. Briley, Robert Sharp, Jessica B. Felix, David D. Moore, Stephanie A. Pangas, Sean M. Hartig

## Abstract

**Objective:** White adipose tissue (WAT) expansion regulates energy balance and overall metabolic homeostasis. WAT absence or loss occurring through lipodystrophy and lipoatrophy contributes to the development of dyslipidemia, hepatic steatosis, and insulin resistance. We previously demonstrated the sole small ubiquitin-like modifier (SUMO) E2-conjuguating enzyme Ubc9 represses human adipocyte differentiation. Germline and other tissue-specific deletions of *Ubc9* frequently cause lethality in mice. As a result, the role of Ubc9 during WAT development remains unknown.

**Methods:** To determine how Ubc9 impacts body composition and energy balance, we generated adipocyte-specific *Ubc9* knockout mice (*Ubc9*^*a-KO*^). CRISPR/Cas9 gene editing inserted loxP sites flanking exons 3 and 4 at the *Ubc9* locus. Subsequent genetic crosses to AdipoQ-Cre transgenic mice allowed deletion of *Ubc9* in white and brown adipocytes. We measured multiple metabolic endpoints that describe energy balance and carbohydrate metabolism in *Ubc9*^*a-KO*^ and littermate controls during postnatal growth.

**Results:** To our surprise, *Ubc9*^*a-KO*^ mice developed hyperinsulinemia and hepatic steatosis. Global energy balance defects emerged from dysfunctional WAT marked by pronounced local inflammation, loss of serum adipokines, hepatomegaly, and near absence of major adipose tissue depots. We observed progressive lipoatrophy that commences in the early adolescent period.

**Conclusions:** Our results demonstrate that *Ubc9* expression in mature adipocytes is essential for maintaining WAT expansion. Deletion of *Ubc9* in fat cells compromised and diminished adipocyte function that provoked WAT inflammation and ectopic lipid accumulation in the liver. Our findings reveal an indispensable role for *Ubc9* during white adipocyte expansion and endocrine control of energy balance.

## 1. INTRODUCTION

White adipocytes safely sequester lipids and protect peripheral metabolic tissues from ectopic lipid accumulation. Consequently, the failure of integral lipid metabolism responses and reduced adipose tissue expandability during chronic states of positive energy balance contribute to the development of insulin resistance, obesity, and fatty liver disease [1]. Similar metabolic abnormalities develop in patients with lipodystrophy where a lack of adipose tissue greatly diminishes storage of surfeit energy and endocrine actions leading to insulin resistance, hepatic steatosis, and dyslipidemia [2]. Therefore, healthy adipose tissue development mediates key aspects of metabolic homeostasis.

Physiologic WAT expansion occurs through both increased adipocyte size (hypertrophy) and number (hyperplasia). White adipocyte differentiation requires a cascade of transcription factors that activate PPARγ, the master regulator of adipocyte differentiation [3]. Adipocyte differentiation requires PPARγ [4] to partner with distinct transcriptional co-regulators that coordinate brown and white adipocyte-specific gene expression [5, 6]. Gene deletion or tissue-specific disruption of PPARγ impairs adipogenesis and results in severe lipodystrophy [4, 7–9]. The mechanisms that enable adipose tissue development carry broad basic research and clinical implications for understanding energy balance disorders.

We previously showed Ubc9 depletion in human subcutaneous preadipocytes accelerates fat cell differentiation [10]. Here, we generated an adipocyte-specific *Ubc9* knockout (*Ubc9*^*a-KO*^) mouse to investigate how Ubc9 regulates whole body energy balance. Surprisingly, WAT mass fails to expand in young male and female *Ubc9*^*a-KO*^ mice. Progressive lipoatrophy and compromised adipocyte function likely provoke WAT inflammation and ectopic lipid accumulation leading to insulin resistance, impaired brown adipose tissue (BAT) function, and intolerance to cold temperatures. Taken together, our findings reveal unexpected roles for *Ubc9* in white adipocyte function and expansion.

## 2. METHODS

### 2.1. Animals

All procedures with animals have been approved by the Institutional Animal Care and Use Committee of Baylor College of Medicine (animal protocol AN-6411). Experimental animals received humane care according to criteria in the “Guide for the Care and Use of Laboratory Animals” (8th edition, revised 2011). Experimental animals were housed (no more than 4 per cage) in a barrier-specific pathogen-free animal facility with 12-h dark-light cycle and free access to water and food. All experiments were conducted using littermate-controlled male and female mice maintained on a C57BL/6J background. At the end of experiments, mice were euthanized by cervical dislocation while under isoflurane anesthesia. After euthanasia, tissues were collected, flash-frozen in liquid N_2_, and stored at −80°C until use. All experiments adhered to ARRIVE Guidelines.

### 2.2. Generation of a Conditional Ubc9 Allele

*Ubc9*^*fl/fl*^ mice were generated by the Genetically Engineered Rodent Models Core at BCM. Two separate and validated single guide RNAs (sgRNAs) and two single-stranded oligonucleotide (ssODN) donors with loxP sequences were used to insert loxP sites 5’ and 3’ of exons 3 and 4 of the mouse *Ube2i* (*Ubc9*) gene, respectively. To minimize the probability of off-target events, only sgRNAs predicted to have off-target sites with three mismatches or more were used to target Cas9 endonuclease activity to intronic sequences flanking exon 3 and 4. Two hundred C57BL/6NJ pronuclear-stage zygotes were co-injected with 100ng/μl Cas9 mRNA, 20ng/μl sgRNA (each), and 100ng/μl of ssODNs [11]. Following micro-injection, zygotes were transferred into pseudopregnant ICR recipient females at approximately 25–32 zygotes per recipient. Sanger sequencing of cloned loxP sites and founder line genotyping from mouse genomic DNA confirmed loxP insertions and sequence fidelity. *Ubc9*^*fl/fl*^ mice were crossed with AdipoQ-Cre (Jackson Laboratory #028020) to generate adipocyte-specific *Ubc9* knockout (*Ubc9*^*a-KO*^) and littermate controls (*Ubc9*^*fl/fl*^).

### 2.3. Genotyping

DNA extracted from mouse ear clips was used in PCR reactions with primers designed to detect the 5’ (P1 - AGGTAGGGGTGGCTTAGAGG, P2 - GGTTCATTGTGCCATCAGGG) and 3’ (P3 - CAAGTCCCAGGGTAGATGCG, P4 - CAGCTCAGACCTGGCCTTAC) loxP sequences and run on agarose gels. AdipoQ-Cre transgenic mice were genotyped according to the protocol provided by the Jackson Laboratory.

### 2.4. Antibodies and Western Blotting

Tissue and whole cell lysates were prepared in Protein Extraction Reagent (Thermo Fisher) supplemented with Halt Protease and Phosphatase Inhibitor Cocktail (Thermo Fisher). Immunoblotting was performed with lysates run on 4-12% Bis-Tris NuPage gels (Life Technologies) and transferred onto Immobilon-P Transfer Membranes (Millipore) followed by antibody incubation. Immunoreactive bands were visualized by chemiluminescence. The following antibodies were used for immunoblotting: α-HSP90 (Cell Signaling #4877), α-Ubc9 (Cell Signaling #4786), α-ADIPOQ (Genetex #GTX112777), and α-Cre (Cell Signaling #7803).

### 2.5. Cell Culture

To validate Cre-inducible deletion of *Ubc9*, fibroblasts were isolated from the inguinal white adipose tissue (iWAT) stromal vascular fraction (SVF) of *Ubc9*^*fl/fl*^ mice. iWAT depots were digested in PBS containing collagenase D (Roche, 1.5 U/ml) and dispase II (Sigma, 2.4 U/ml) supplemented with 10 mM CaCl_2_ at 37°C for 45 min. The primary cells were filtered twice through 70 μm strainers and centrifuged to collect the SVF. For adenovirus transduction, SVF cells were incubated with adenoviral Cre recombinase or green fluorescent protein (GFP) in DMEM/F12 medium containing Glutamax (ThermoFisher) and 10% fetal bovine serum for 24 hours. After replacing the medium once, cells were cultured for 48 hours before lysate preparation. Adenovirus expressing Cre recombinase or GFP were provided by the BCM Gene Vector Core.

### 2.6. Indirect Calorimetry

*Ubc9*^*a-KO*^ and littermate controls (*Ubc9^fl/fl^)* were maintained on normal chow and housed at room temperature in Comprehensive Lab Animal Monitoring System (CLAMS) home cages (Columbus Instruments). Oxygen consumption, CO_2_ emission, energy expenditure, food and water intake, and activity were measured for 6 days (BCM Mouse Metabolic and Phenotyping Core). Mouse body weight was recorded, and body composition examined by MRI (Echo Medical Systems) prior to indirect calorimetry. Statistical analysis of energy balance was performed by ANCOVA with lean body mass as a co-variate using the CalR web-based tool [12].

### 2.7. Glucose and Insulin Tolerance Tests

To determine glucose tolerance, mice were fasted for 16 hours and glucose was administered (1.5 g/kg body weight) by intraperitoneal (IP) injection. To determine insulin tolerance, mice were fasted four hours prior to insulin IP injection (1.5 U/kg body weight). Blood glucose levels were measured by handheld glucometer.

### 2.8. ELISAs and Lipid Assays

Fed serum levels were used to measure insulin (#EZRMI-13K; Millipore), leptin (#90030; Crystal Chem), adiponectin (#KMP0041; Thermo Fisher), free fatty acids (#sfa-1; ZenBio), and FGF21 (#MF2100; R&D Systems). Hepatic triglyceride content was quantified by Thermo Scientific Triglycerides Reagent (#TR22421) and normalized per gram of liver tissue.

### 2.9. Histology

Formalin-fixed paraffin-embedded adipose and liver tissue sections were stained with hematoxylin and eosin (H/E) by the BCM Human Tissue Acquisition and Pathology Core. Images were captured using a Nikon Ci-L Brightfield microscope.

### 2.10. qPCR

Total RNA was extracted using the Direct-zol RNA MiniPrep kit (Zymo Research). cDNA was synthesized using iScript (Bio-Rad). Relative mRNA expression was measured with SsoAdvanced Universal Probes Supermix reactions read out with a QuantStudio 3 real-time PCR system (Applied Biosystems). TATA-box binding protein (*Tbp)* was the invariant control. Roche Universal Probe Gene Expression Assays were used as previously described [13].

### 2.11. Cold Tolerance Test

Six-month old male *Ubc9*^*fl/fl*^ and *Ubc9*^*a-KO*^ mice were individually housed with water and exposed to cold temperature (4°C) for 2.5 hours. A temperature probe was placed subcutaneously on top of the intrascapular BAT two days prior to the cold tolerance test. Temperature recordings were measured in duplicate at room temperature (time=0) and during cold exposure every 30 min.

### 2.12. Statistical Analyses

Statistical significance was assessed by unpaired Student’s t-test. All data are presented as mean ± SEM, unless otherwise stated. Our primary threshold for statistical significance was p<0.05.

## 3. RESULTS

### 3.1. Generation of conditional *Ubc9* knockout mice

*Ubc9* is expressed ubiquitously across all mouse tissues [14] and knockout strategies cause lethality and sterility [14–17]. To this end, we used CRISPR/Cas9 gene editing to generate a floxed *Ubc9* allele (*Ubc9*^*fl/fl*^) to explore tissue-specific roles for *Ubc9*. LoxP sites flanking exons 3 and 4 of the *Ubc9* locus were introduced by homology-directed repair **(Figure 1A)** and the *in vivo* presence of loxP sites in the targeted regions was confirmed by genotyping of potential founder mice **(Figure 1B)**. We verified that the loxP sites targeted the *Ubc9* locus by transfecting *Ubc9*^*fl/fl*^ SVF with adenovirus expressing Cre recombinase. Immunoblot analysis of whole cell lysates demonstrated near total deletion of Ubc9 protein levels following Cre recombination compared to adenovirus GFP transductions **(Figure 1C)**. To study the effects of *Ubc9* deletion specifically in mature fat cells, we generated adipose-specific *Ubc9* knockout mice (*Ubc9*^*a-KO*^) by crossing *Ubc9*^*fl/fl*^ animals with AdipoQ-Cre transgenic mice **(Figure 1D)**. Reproductive fitness and female nursing were unaffected by *Ubc9*^*a-KO*^ and all pups were viable and born at the expected Mendelian ratio. PCR analysis demonstrated Cre recombination of the *Ubc9* locus generated a deletion product (red arrow) in the gonadal WAT (gWAT) from *Ubc9*^*a-KO*^ mice that was absent in *Ubc9*^*fl/fl*^ controls **(Figure 1E)**. The full length, unrecombined product (green arrow) was also detected in *Ubc9*^*a-KO*^, but at lower levels than control, suggesting contributions of Cre-negative cells in the SVF to the PCR product. Similarly, Ubc9 protein was reduced in inguinal WAT (iWAT) and BAT compared to *Ubc9*^*fl/fl*^ controls **(Figure 1F)**. Ubc9 loss in WAT did not affect body weight or weight gain in males or females **(Figure 1G)**.

**Figure 1.**
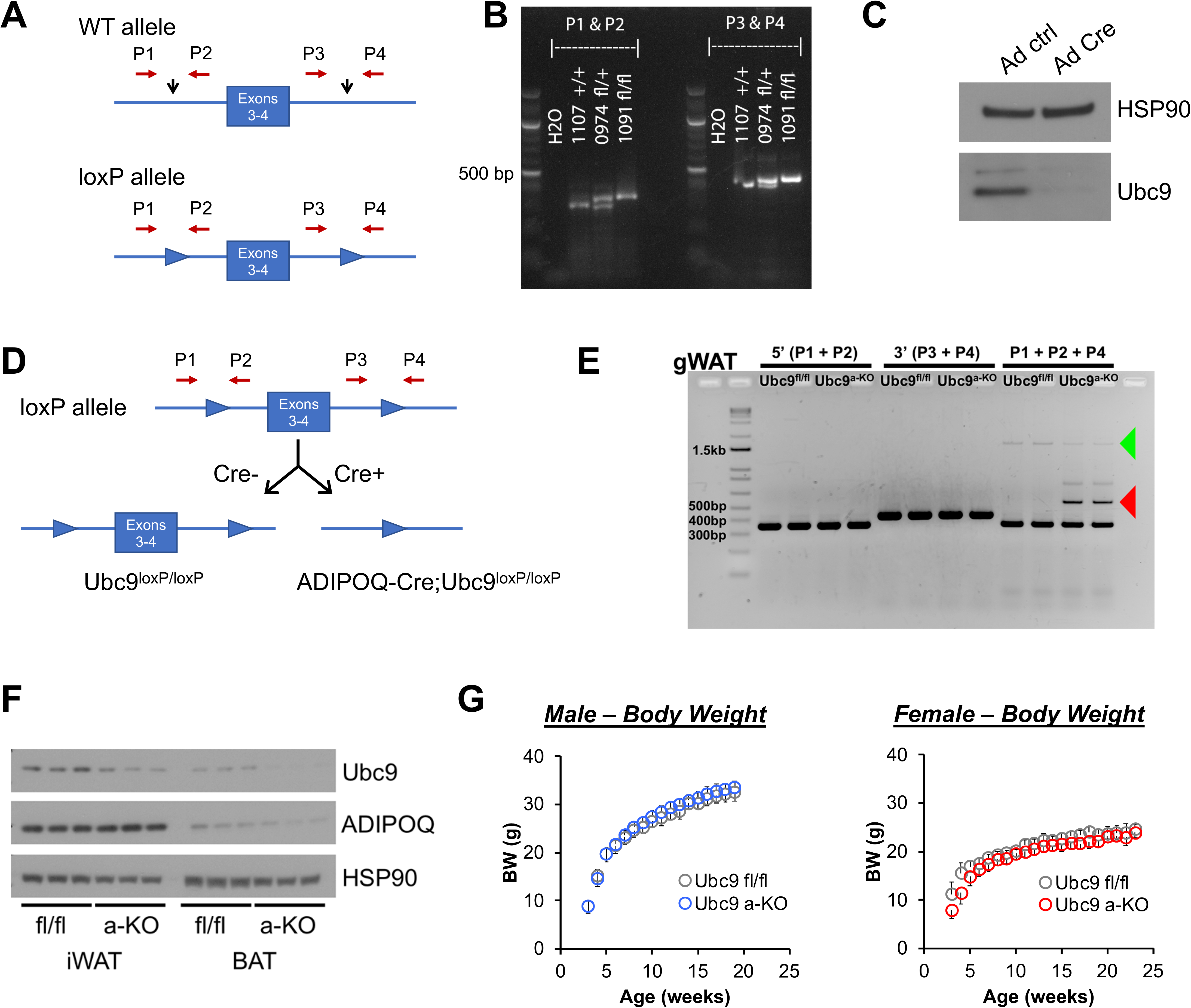
Generation of conditional *Ubc9* gene deletion mice. **(A)** *Ubc9*^*fl/fl*^ mice were generated using CRISPR/Cas9 gene editing. Exons 3 and 4 of the *Ube2i* (*Ubc9*) gene were targeted by sgRNAs designed complementary to intronic sequences flanking the exons, then loxP sequences were introduced by DNA donor oligonucleotides. LoxP sites were inserted before exon 3 and after exon 4 (black arrows). Primers detect the 5’ (P1, P2) and 3’ (P3, P4) loxP sequences. **(B)** PCR analyses of floxed alleles at the targeted loci in genomic DNA extracted from ear clips of wild-type (+/+), fl/+, and fl/fl mice. PCR products were run on agarose gels with expected band sizes for P1-P2: wild-type (+) 319 bp and loxP allele (fl) 353 bp and P3-P4: wild-type (+) 427 bp and loxP allele (fl) 461 bp. The image shows a wild-type control #1107 +/+, founder heterozygous mouse #974 fl/+, and homozygous F2 offspring #1091 fl/fl. **(C)** Western blot analysis of Ubc9 expression in iWAT SVF cells from *Ubc9*^*fl/fl*^ mice after transduction with adenoviral GFP or Cre. **(D)** Strategy for generating adipocyte-specific *Ubc9* gene deletion. *Ubc9*^*fl/fl*^ mice were bred with AdipoQ-Cre mice to generate adipocyte-specific *Ubc9* knockout (*Ubc9*^*a-KO*^) and *Ubc9*^*fl/fl*^ (control) mice. **(E)** To validate deletion of *Ubc9*, genomic DNA was extracted from *Ubc9*^*a-KO*^ and *Ubc9*^*fl/fl*^ gWAT samples and PCR products were run on an agarose gel to detect the 5’ (P1, P2 primers) and 3’ (P3, P4 primers) loxP sequences, as well as a product that spans exons 3-4 (P1+P2+P4; 1597 bp, green arrow) or the deletion product (509 bp, red arrow). **(F)** Western blots of whole tissue lysates from iWAT and BAT of seven-day old mice were probed for the indicated proteins. **(G)** *Ubc9*^*a-KO*^ and *Ubc9*^*fl/fl*^ (control) mice were weighed for up to 23 weeks in male (blue/gray; n=12-14, mean +/− SD) and female (red/gray; n=6-8, mean +/− SD).

### 3.2. Hepatosteatosis and insulin resistance in adult *Ubc9*^*a-KO*^ mice

Despite equivalent body weights, six-month old male and female *Ubc9*^*a-KO*^ mice developed fatty liver disease on a normal chow diet. Both male and female *Ubc9*^*a-KO*^ mice displayed hepatic lipid droplet accumulation **(Figure 2A)** with significantly increased triglyceride content **(Figure 2B)**. FGF21 levels were increased 4-fold in *Ubc9*^*a-KO*^ serum compared to controls **(Figure 2C)**, suggesting elevated stress responses in the liver [18]. *Ad libitum* fed glucose levels trended higher in *Ubc9*^*a-KO*^ **(Figure 2D)**, despite dramatically elevated levels of serum insulin **(Figure 2E)**, which reflected a state of insulin resistance noted by higher HOMA-IR values **(Figure 2F)**. Interestingly, we detected ~14-fold higher serum insulin levels in male *Ubc9*^*a-KO*^ mice while females exhibited only a 4-fold increase compared to controls. To test the consequences of dyslipidemia on glucose homeostasis, we performed insulin **(Figure 2G)** and glucose tolerance tests **(Figure 2H)**. *Ubc9*^*a-KO*^ male and female mice were more insulin resistant relative to controls. Interestingly, male *Ubc9*^*a-KO*^ showed improved glucose intolerance with a lower fasting glucose level. In contrast, fasting glucose was elevated in female *Ubc9*^*a-KO*^ compared to control, with nominal effects on overall glucose tolerance. This sex discrepancy in glucose tolerance can be explained by higher amounts of insulin in male *Ubc9*^*a-KO*^ mice that overcome the glucose challenge.

**Figure 2.**
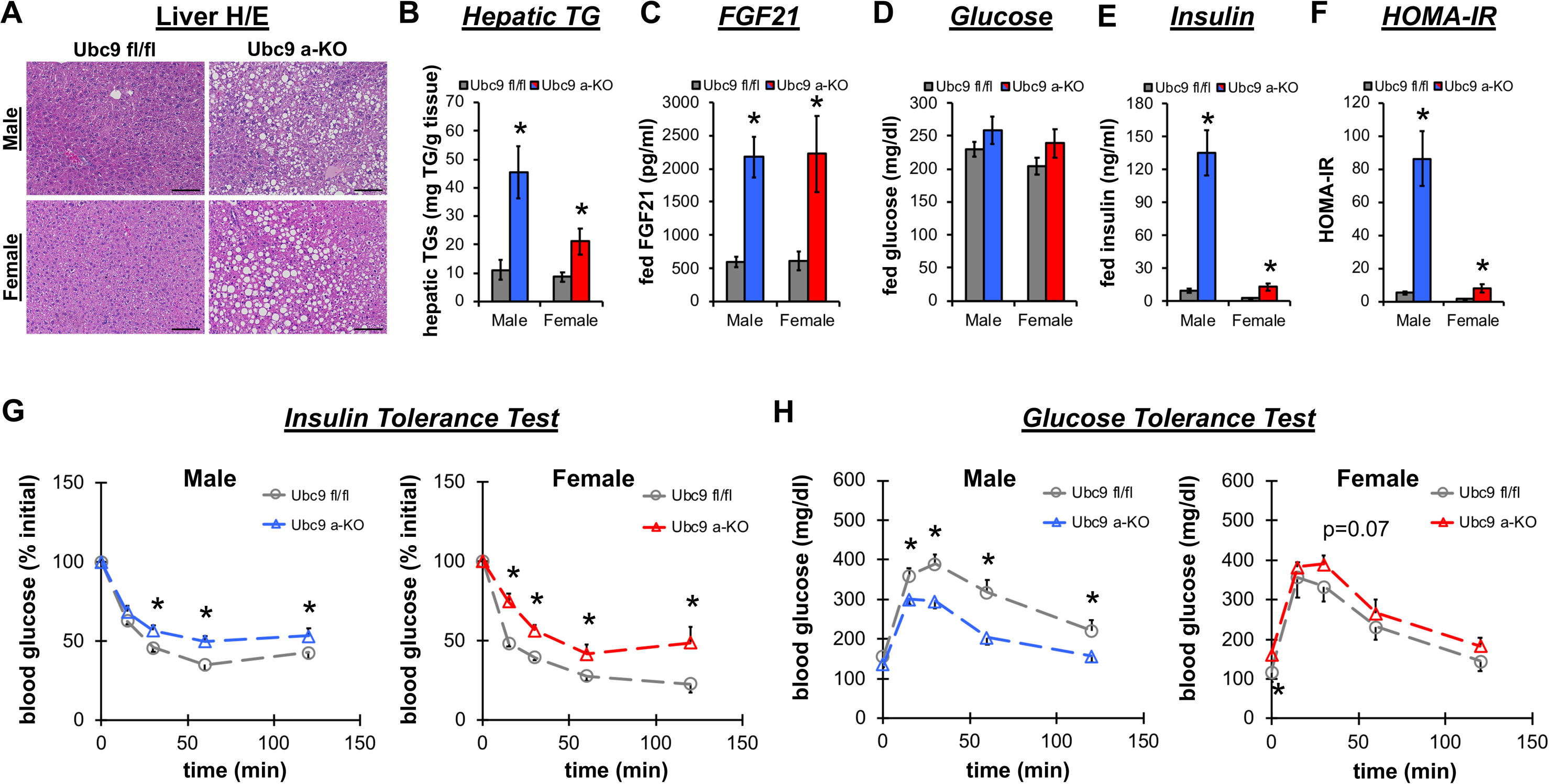
Adipocyte-specific *Ubc9* knockout mice develop insulin resistance and hepatic steatosis. **(A)** Representative H/E stained liver sections from male (top row) and female (bottom row) *Ubc9*^*a-KO*^ and *Ubc9*^*fl/fl*^ control mice show lipid droplet accumulation in *Ubc9*^*a-KO*^ mice. Scale bar 100 μm. **(B)** Hepatic triglycerides (TGs) per gram liver tissue (n=7/group). **(C)** Fed serum FGF21 (pg/ml) (male, blue n=11-12; female, red n=7-9). **(D)** Fed serum glucose (mg/dl), **(E)** insulin (ng/ml), and **(F)** HOMA-IR measurements (male, blue n=11-12; female, red n=7-9). **(G)** Insulin and **(H)** glucose tolerances tests were performed on *Ubc9*^*fl/fl*^ and *Ubc9*^*a-KO*^ male (blue; n=11-15) and female (red; n=5-7) mice. Data represent mean +/− SEM; *p<0.05. Tissue and serum chemistry analyses were performed in six-month old adult mice.

### 3.3. Adipocyte-specific *Ubc9* deletion increases energy expenditure

We next performed a series of experiments to determine the effects of *Ubc9*^*a-KO*^ on energy expenditure and BAT thermogenic functions. Adult male *Ubc9*^*a-KO*^ mice were placed in CLAMS home cages to assess heat production (energy expenditure) by indirect calorimetry. *Ubc9*^*a-KO*^ mice exhibited increased energy expenditure **(Figure 3A)** and primarily used glucose as a fuel source during the light period **(Figure 3B)** as indicated by elevated respiratory exchange ratio (RER). The genotype effects were significant when lean body mass was used as the covariate. Poor metabolic flexibility coupled with increased energy expenditure likely motivated greater food intake in *Ubc9*^*a-KO*^ mice **(Figure 3C)**. Increased energy expenditure was not associated with higher activity levels **(Figure 3D)**. Histological assessment of H/E stained BAT sections from *Ubc9*^*a-KO*^ mice demonstrated lipid accumulation in large unilocular fat droplets, in contrast to the small multi-locular fat droplets characteristic of controls **(Figure 3E)**. *Ubc9* knockout in adipose tissues caused dysfunctional BAT gene expression as reflected by reduced levels **(Figure 3F)** of adipocyte markers (*AdipoQ*, *Pparg2*, *Pdrm16*) and key lipid metabolism genes (*Ucp1*, *Cidea*, *Dio2*, *Pgc1a*, *Acaca*, *Acadm*, *Acadl*). To test the functional output of BAT, we performed a cold tolerance test in the absence of food. Before cold exposure, BAT temperature was reduced in *Ubc9*^*a-KO*^ mice compared to controls and dramatically dropped after 2.5 hours at 4°C **(Figure 3G)**, demonstrating an inability of *Ubc9*^*a-KO*^ mice to defend body temperature. Collectively, these experiments demonstrate *Ubc9*^*a-KO*^ causes a hypermetabolic phenotype coupled with metabolic inflexibility and BAT thermogenic defects.

**Figure 3.**
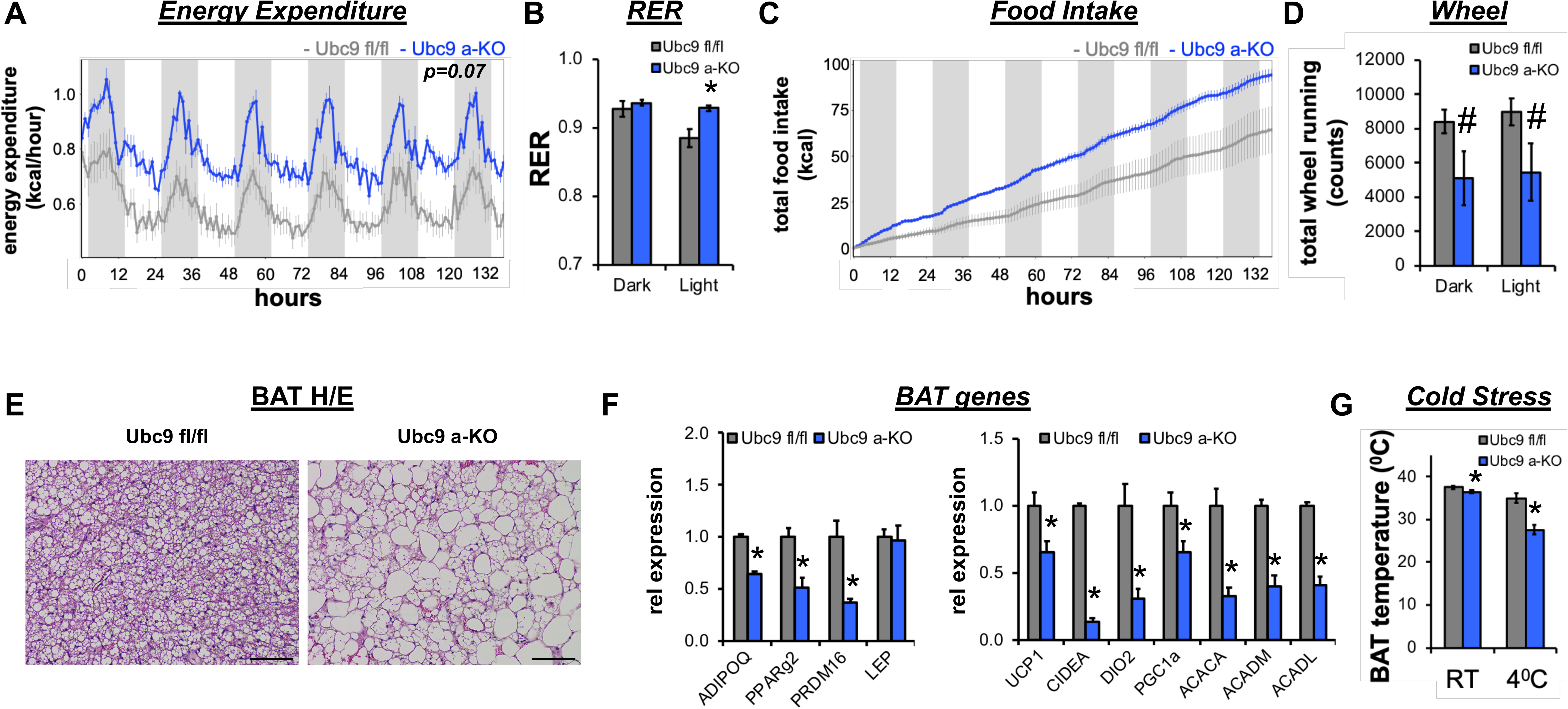
Energy balance defects caused by adipocyte-specific *Ubc9* deletion. Six-month old *Ubc9*^*fl/fl*^ (gray) and *Ubc9*^*a-KO*^ (blue) male mice were individually housed and monitored in CLAMS home cages for 6 days (n=4-5). Recorded traces of **(A)** energy expenditure (kcal/hour), **(B)** averaged RER during dark and light periods, and **(C)** cumulative food intake. Statistical analysis of energy balance was performed by ANCOVA with lean body mass as a co-variate. **(D)** Activity was measured by wheel running. **(E)** H/E stained BAT sections from male *Ubc9*^*fl/fl*^ and *Ubc9*^*a-KO*^ mice show large unilocular lipid droplets in *Ubc9*^*a-KO*^ mice. Scale bar 100 μm. **(F)** Relative gene expression by qPCR for markers of BAT function (n=4/group). **(G)** Temperature probes were inserted under the skin to monitor intrascapular BAT temperature before (room temperature, RT) and after 2.5 hours of cold (4°C) exposure (n=3-5). Data represent mean +/− SEM; *p<0.05, #p<0.10.

### 3.4. Adipocyte-specific *Ubc9* deletion increases WAT inflammation and stress

To assess the morphological changes associated with *Ubc9* knockout in adipose tissues, we performed H/E staining of gWAT and iWAT tissue sections in adult male and female mice. Pronounced immune cell infiltration was observed in gWAT of *Ubc9*^*a-KO*^ mice **(Figure 4A)**, distinct from the commonly described crown-like structures typically associated with gWAT [19]. Similarly, we observed dispersed immune cell accumulation amongst stromal cells and large adipocytes in the iWAT of *Ubc9*^*a-KO*^ mice. Primary WAT depots from both sexes showed marked stroma invasion and few mature adipocytes. Consistent with reduced numbers of adipocytes, serum levels of adiponectin, leptin, and free fatty acids were significantly reduced in *Ubc9*^*a-KO*^ mice compared to controls **(Figure 4B)**. Adipocyte-specific deletion of *Ubc9* reduced hallmark adipocyte genes (*AdipoQ*, *Pparg2*, *Lep*, *Fabp4*) in WAT associated with upregulation of immune cell markers (*F4/80*, *Foxp3*, *Tnfa*, *Ifng*) and collagen (*Col1a1*) in both the gWAT **(Figure 4C)** and iWAT (**Figure 4D)**. These data suggest impaired fat storage and WAT expansion in *Ubc9*^*a-KO*^ mice induces stress and inflammatory responses.

**Figure 4.**
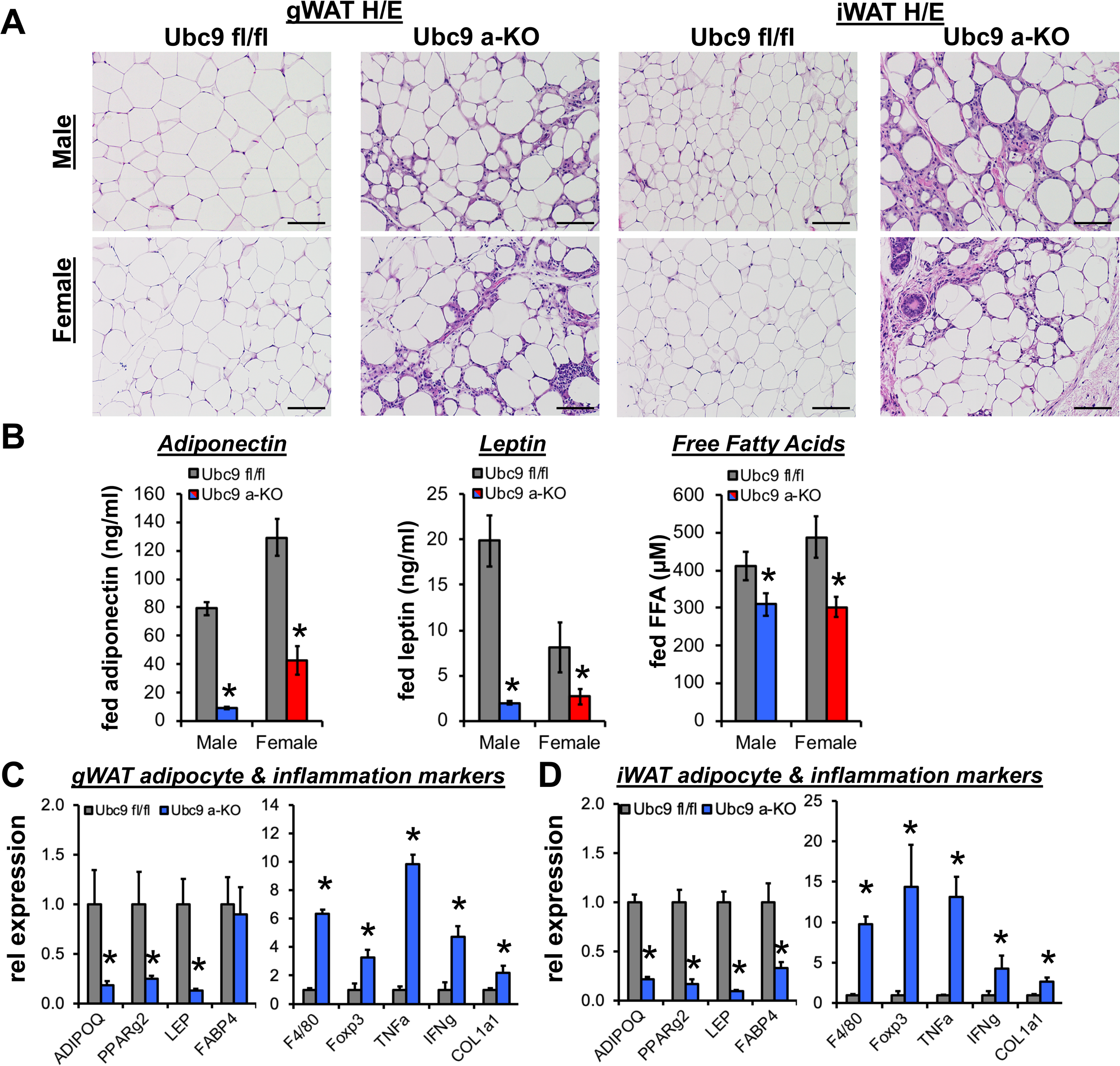
*Ubc9*^*a-KO*^ mice display impaired WAT function. **(A)** H/E stained sections from gonadal and inguinal WAT from male (top row) and female (bottom row) mice show substantial stromal cell infiltration in WAT of *Ubc9*^*a-KO*^ mice. Scale bar 100 μm. **(B)** Fed serum adiponectin (ng/ml), leptin (ng/ml), and free fatty acids (μM; FFA) in male (n=11-12) and female (n=7-9) *Ubc9*^*a-KO*^ mice compared to *Ubc9*^*fl/fl*^ controls. Relative gene expression by qPCR (n=4/group) for markers of mature adipocytes and inflammation in **(C)** gonadal and **(D)** inguinal WAT of male mice. Data represent mean +/− SEM; *p<0.05. All analyses were performed in six-month old adult mice.

### 3.5. Adipocyte-specific *Ubc9* deletion impairs WAT expansion

The inability of WAT depots to safely sequester lipids causes ectopic accumulation of energy in peripheral organs and lipodystrophic phenotypes [20]. Magnetic resonance imaging demonstrated reduced fat mass and increased lean mass in six-month old *Ubc9*^*a-KO*^ male and female mice compared to controls **(Figure 5A)**. Further examination of tissue weights revealed grossly visible reductions (−80%) in iWAT and gWAT depots in male and female *Ubc9*^*a-KO*^ mice **(Figure 5B)**. Reduced fat storage in primary WAT depots resulted in significantly higher liver weights associated with gross morphological changes confirming ectopic lipid accumulation and the previously observed hepatic steatosis. Similarly, “whitening” of BAT was apparent at necropsy, with increased BAT weight detected in female *Ubc9*^*a-KO*^ mice **(Figure 5C)**. Examining body weight **(Figure 5D)** and WAT mass from seven days to six months of age revealed progressive expansion of WAT mass in *Ubc9*^*fl/fl*^ control mice, as expected **(Figure 5E)**. In contrast, *Ubc9*^*a-KO*^ mice failed to expand WAT depots beginning at two months of age and by six months, WAT mass almost completely disappeared relative to *Ubc9*^*fl/fl*^ controls. Collectively, these data demonstrate adipocyte-specific deletion of Ubc9 impairs WAT expansion leading to ectopic lipid accumulation.

**Figure 5.**
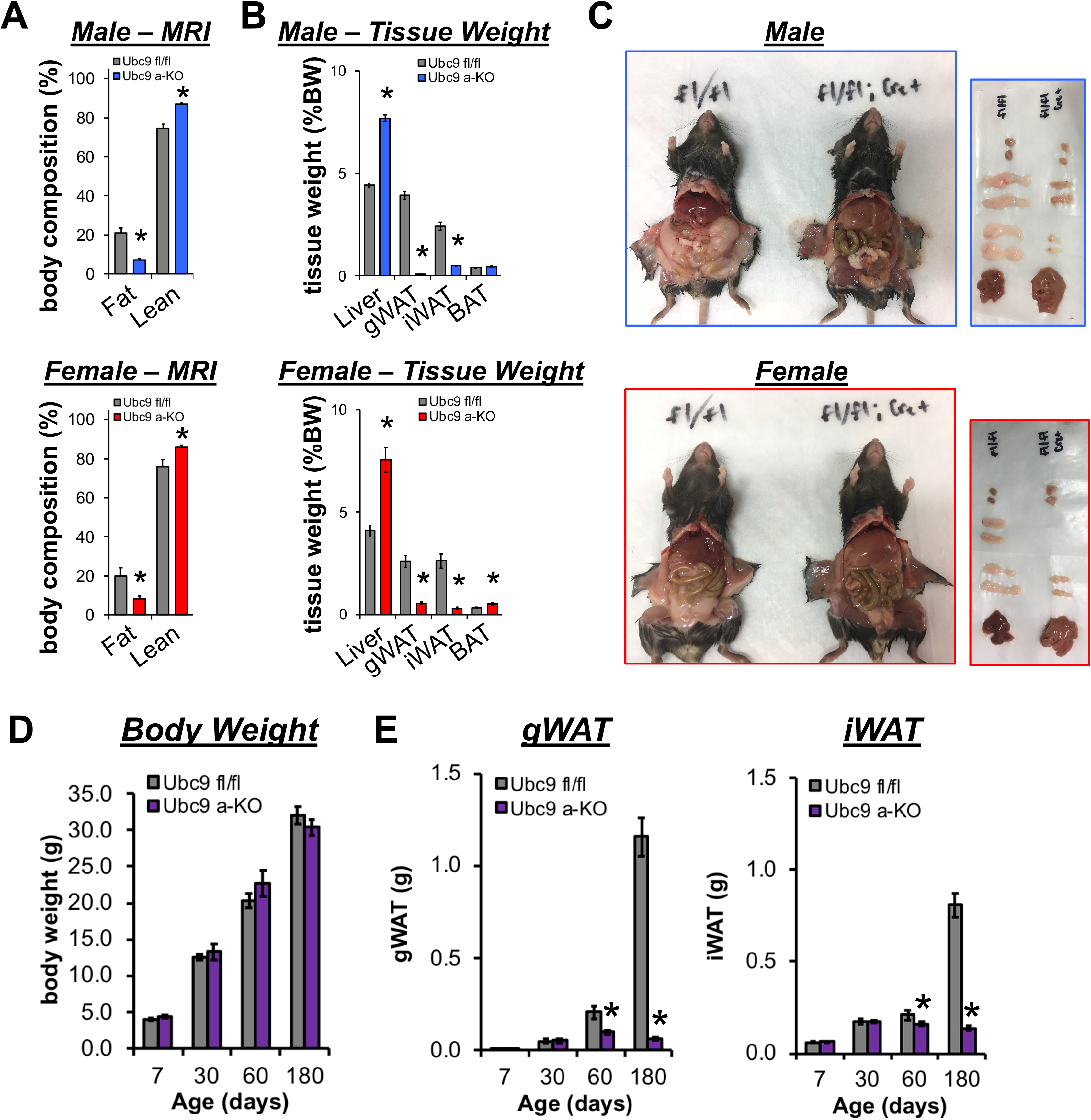
Progressive WAT loss in *Ubc9*^*a-KO*^ mice during postnatal growth. **(A)** Assessment of fat and lean mass (% body weight; male n=11-15, female n=5-7) with **(B)** corresponding tissues weights (% body weight; male n=15-16, female n=8) and **(C)** necropsy images from six-month old *Ubc9*^*fl/fl*^ and *Ubc9*^*a-KO*^ mice. Images of excised tissues demonstrate gross morphological increases in liver size, reductions in iWAT and gWAT, and lighter coloring of the liver and BAT. **(D)** Body weight and **(E)** WAT weights (g) in male and female mice at 7 (n=5-9), 30 (n=4/group), 60 (n=4/group), and 180 (n=23-24/group) days of age. Data represent mean +/− SEM; *p<0.05.

## 4. DISCUSSION

Our previous work identified the E2 SUMO conjugating enzyme, Ubc9, as a negative regulator of fat cell differentiation in human subcutaneous preadipocytes [10]. Here, we generated adipocyte-specific *Ubc9* knockout (*Ubc9*^*a-KO*^) mice to define the *in vivo* functions of Ubc9 in mature fat cells. To our surprise, male and female *Ubc9*^*a-KO*^ mice developed a hypermetabolic phenotype and progressive WAT lipoatrophy associated with immune cell infiltration and decreased circulating adipokines. *Ubc9*^*a-KO*^ mice exhibited additional hallmark features of lipodystrophy, including ectopic lipid deposition in the liver, insulin resistance, and metabolic inflexibility. These studies demonstrate previously unrealized functions of Ubc9 in mouse WAT that are essential for mature adipocyte function and survival.

Patients with congenital general (CGL) or familial partial (FPLD) lipodystrophy exhibit profound insulin resistance, hepatic steatosis, and dyslipidemia. Female patients in particular develop more severe metabolic complications than males [21, 22]. Similarly, impaired glucose and insulin tolerance in *Ubc9*^*a-KO*^ mice were more pronounced in females, attributed in part to a much lower hyperinsulinemic response compared to males. In terms of fat mass, CGL patients have extreme deficits at birth due to autosomal recessive mutations, while FPLD patients develop progressive fat loss beginning at puberty [2, 21–23]. *Ubc9*^*a-KO*^ mice resemble FPLD because WAT mass was normal in *Ubc9*^*a-KO*^ mice after birth, but began to decrease during puberty, with as little as 2% remaining by six months. The timing of fat loss in *Ubc9*^*a-KO*^ mice is consistent with a period of hypertrophic WAT expansion [24], impairments of which degrade adipocyte maturation and endocrine functions. Accordingly, inhibition of SUMOylation in differentiated cells [25] decreased adipocyte lipid storage and maturation (*Pparg*, *C/ebpa*, *Fabp4*). Collectively, these observations suggest Ubc9 has a distinct role in adult adipocyte function, maturation, and survival.

A few lipodystrophic mouse models develop progressive lipoatrophy and hepatic steatosis, such as adipocyte-specific knockout of Akt1/Akt2 [26], IR [27], Raptor [28], and PPARγ [4]. It is worth noting that fed glucose levels remained normal in *Ubc9*^*a-KO*^ mice despite hyperinsulinemia. This observation suggests beta cell secretion partly corrects the mild insulin resistance in *Ubc9*^*a-KO*^ mice. Our model resembles many features of progressive lipoatrophy observed in PPARγ knockout models [4, 8]. Lipoatrophy in fat-specific PPARγ knockout mice occurs because adipocytes lost lipids and shrank, consistent with the pivotal roles of PPARγ as a master regulator of adipose tissue formation [4, 29]. PPARγ ablation in Sox2-expressing cells (PPARγ^Δ/Δ^ skirts embryonic lethality and mice survive without WAT and BAT [8]. PPARγ ^Δ/Δ^ show a hypermetabolic phenotype accompanied by higher energy expenditure and hyperphagia. Similar to *Ubc9*^*a-KO*^ mice, PPARγ^Δ/Δ^ mice show higher RER and glucose oxidation that derives from more lean mass. The liver dominates mass-specific metabolic rates [30] and, for this reason, the hepatomegaly and hyperinsulinemia present in *Ubc9*^*a-KO*^ likely contribute to greater glucose oxidation and metabolic inflexibility. Our observations expand on these studies and document a putative role for Ubc9 and PPARγ interactions [10] in maintaining adipose tissue homeostasis during postnatal growth.

For reasons that remain unclear, *Ubc9* expression peaks early in differentiation of 3T3-L1 adipocytes, when adiponectin remains low, to regulate expression of the key adipogenic transcription factors PPARγ, C/EBPα, and C/EBPβ [31]. Adiponectin expression occurs late in adipogenesis [32] and AdipoQ-Cre mediated ablation of *Ubc9* occurs in relatively mature adipocytes. While impaired adipogenesis underlies development of lipodystrophy in some models [8, 33–35], it is unlikely to account for lipoatrophy *in Ubc9^a-KO^* mice. The more dramatic lipodystrophy in older *Ubc9*^*a-KO*^ likely results from apoptosis and inflammation, consistent with the pivotal roles of *Ubc9* in cell survival across many tissue types [14]. Cell survival also depends on preservation of nuclear architecture, evident by the embryonic lethality of Ubc9 deficiency [16]. Additionally, Ubc9 occupies transcriptional start sites of functional genes integral for cell growth and proliferation [36]. However, loss of *Ubc9* induces cell growth arrest, due in part, to impaired chromosome segregation, which contributes to reduced cell viability [36]. Along these lines, it will now be important to use mass spectrometry and lineage tracing methods to identify the most critical metabolic and transcription factors downstream of Ubc9 functions that explain why Ubc9 allows WAT expansion during aging.

Ubc9 depletion provokes senescence in fibroblasts [36]. These data suggest Ubc9 maintains a balance between cell proliferation and terminal differentiation. Premature senescence and growth arrest in young mice when fat pads require rapid adaptation to nutritional demands may compromise WAT function and expansion. In turn, this would be expected to accelerate differentiation of a limited pool of adipocyte progenitors [37–39]. This concept is consistent with observations in the small intestine of adult *Ubc9* knockout mice whereby elevated proliferation and apoptosis associates with rapid depletion of stem cells [14]. Clearing senescent cells reverses fat loss associated with aging and restores expression of *Pparg*, *C/ebpa*, and other mature adipocyte markers [40, 41]. Thus, *Ubc9*^*a-KO*^ may induce premature aging resulting from growth arrest and senescence associated with dysfunctional preadipocyte turnover.

Conditions of lipodystrophy and pathologic obesity suffer from a restricted capacity to expand white adipose depots to meet the demand for nutrient storage, which drives adipocyte dysfunction and WAT inflammation, along with ectopic lipid accumulation and insulin resistance [42]. Ultimately, identifying factors that enable healthy WAT expansion carries significant implications for treating diseases associated with fat storage and metabolism, including obesity and lipodystrophy. Our data reveal Ubc9 exerts an essential role in lipid storage and WAT expansion of young adult mice. Consequently, adipocyte-specific loss of Ubc9 impairs WAT expansion, contributing to lipoatrophy through elevated WAT inflammation and adipocyte loss.

The striking phenotype of *Ubc9*^*a-KO*^ mice reveal the cell-autonomous necessity of Ubc9 in healthy adipose tissue maintenance and whole-body energy balance. As a rate-limiting E2 SUMO conjugating enzyme necessary for SUMOylation, this mouse model provides a valuable tool for understanding how the low-abundance post-translational modification SUMOylation [43] broadly affects mature adipocyte function and, more broadly, tissue development. In addition*, Ubc9^a-KO^* mice add to a small list of mouse models to study systemic effects of WAT loss during postnatal development and fundamental aspects of energy balance.

## Abbreviations

a-KO: adipocyte-specific knockout
BAT: brown adipose tissue
CGL: congenital generalized lipodystrophy
CLAMS: Comprehensive Lab Animal Monitoring System
CRISPR: Clustered Regularly Interspace Short Palindromic Repeats
FFA: free fatty acids
FPLD: familial partial lipodystrophy
GFP: green fluorescent protein
gWAT: gonadal white adipose tissue
H/E: Hematoxylin and Eosin
HOMA-IR: Homeostatic Model Assessment of Insulin Resistance
iWAT: inguinal white adipose tissue
loxP: locus of X-over P1
PPARγ: Peroxisome proliferator-activated receptor gamma
RER: respiratory exchange ration
sgRNA: single guide RNA
ssODN: single stranded oligonucleotide
SUMO: Small Ubiquitin-like Modifier
SVF: stromal vascular fraction
TG: triglyceride
Ubc9: Ubiquitin carrier protein 9
WAT: White adipose tissue

## Funding

This work was funded by American Diabetes Association #1-18-IBS-105 and NIH R01DK114356 (to S.M.H.), R01 HD085994 and T32 HD098068 (to S.A.P). This study was also funded (in part) by an award from the Baylor College of Medicine Nutrition and Obesity Pilot and Feasibility Fund (to A.R.C.). This study was also supported, in part, by the Assistant Secretary of Defense for Health Affairs endorsed by the DOD PRMRP Discovery Award (No. W81XWH-18-1-0126 to K.H.K.). This project was supported in part by PHS grant P30DK056338. Core services at BCM utilized in this project were supported with funding from NCI P30-CA125123: Genetically Engineered Rodent Models Core, Human Tissue Acquisition and Histology Core, and the Integrated Microscopy Core.

## Author Contributions

A.R.C. and S.M.H. conceptualized the study. P.M., N.C., A.R.C., and S.M.H. designed experiments. A.R.C. and S.M.H. wrote the manuscript with editorial input from all authors. S.M.H. and A.R.C performed all experiments with assistance as noted: P.K.S. assisted with mouse phenotyping; A.R.C. performed qPCR analysis with assistance from J.B.F., N.C. and R.S.; D.D.M., S.M.B., and S.A.P. performed genotyping and troubleshooting; K.H.K. performed analysis of liver lipids. All work was performed under the supervision of S.M.H.

## Conflicts of Interest

The authors have declared no conflict of interest exists.

## Conflict of Interest Statement

The authors have declared no conflict of interest exists.

## Notes

### Competing Interest Statement

The authors have declared no competing interest.

